# Thermoregulatory Dynamics Reveal Sex-Specific Inflammatory Responses to Experimental Autoimmune Encephalomyelitis in Mice: Implications for Multiple Sclerosis-Induced Fatigue in Females

**DOI:** 10.1101/2022.03.16.484095

**Authors:** Jamshid Faraji, Dennis Bettenson, Stella Babatunde, Tabitha Gangur-Powell, Voon Wee Yong, Gerlinde A.S. Metz

## Abstract

The course of multiple sclerosis (MS) is characterized by striking sex differences in symptoms such as fatigue and impaired thermal regulation, which are associated with aggravated systemic pro-inflammatory processes. The purpose of this study was to replicate these symptoms in experimental autoimmune encephalomyelitis (EAE) in C57BL/6 mice in the quest to advance the preclinical study of non-motor symptoms of MS. Male and female C57BL/6 mice exposed to a mild form of EAE were evaluated for the progression of clinical, behavioural, thermal, and inflammatory processes. We show higher susceptibility in females to EAE than males based on greater clinical score and cumulative disease index (CDI), fatigue-like and anxiety-like behaviours. Accordingly, infrared (IR) thermography indicated higher cutaneous temperature in females from post-induction days 12 to 23. Females also responded to EAE with greater splenic and adrenal gland weights than males as well as sex-specific changes in pro- and anti-inflammatory cytokines. These findings provide the first evidence of a sex-specific thermal response to immune-mediated demyelination, thus proposing a non-invasive assessment approach of the psychophysiological dynamics in EAE mice. The results are discussed in relation to the thermoregulatory correlates of fatigue and how endogenously elevated body temperature without direct heat exposure may be linked to psychomotor inhibition in patients with MS.

## Introduction

Much of the variability in responses to neurological disorders can be explained by gender and/or sex. In humans, it has been known for decades that females are more susceptible than males to multiple sclerosis (MS) (Doss et al., 2021), a chronic immune-mediated demyelinating disease which is characterized by neurodegeneration and neurological disability. Women develop MS 2– 3 times more commonly than men (Doss et al., 2021; Greer & McCombe, 2011; Voskuhl & Palaszynski, 2001). Heightened vulnerability among females has also been observed in preclinical rodent models of MS, such as experimental autoimmune encephalomyelitis (EAE) in mice (Cruz-Orengo et al., 2014; Doss et al., 2021; Hoghooghi et al., 2020; Wiedrick et al., 2021). EAE is a widely used animal model because it involves both the adaptive an innate immune responses (Berard, Wolak, Fournier, & David, 2010; Gerrard et al., 2017), and the formation of perivascular white-matter lesions that are characteristic for human MS (Wiedrick et al., 2021; Wuerch, Mishra, Melo, Ebacher, & Yong, 2022). Like MS, EAE also elicits acute, chronic, and progressive motor disabilities in rodents (Berard et al., 2010; Jensen et al., 2018; Reich, Lucchinetti, & Calabresi, 2018).

While female of many mouse strains are more susceptible to the development of EAE than their male counterparts (Papenfuss et al., 2004), the underlying mechanisms are not fully understood. Sex differences in MS and EAE susceptibility and symptom severity have been attributed to heritability of risk genes and chromosomal influences, sexual dimorphisms in brain anatomy, or sex hormones that influence the production of inflammatory cytokines (Cruz-Orengo et al., 2014; Hoghooghi et al., 2020; Smith-Bouvier et al., 2008; Voskuhl & Palaszynski, 2001; Wilcoxen, Kirkman, Dowdell, & Stohlman, 2000). In turn, the higher female-to-male ratio reflected by inflammatory cytokine responses to EAE (Golden & Voskuhl, 2017; Hoghooghi et al., 2020) may also explain sex disparities in behavioural responses to disease pathogenesis.

In humans, chronic fatigue syndrome (CFS) represents a common clinical sign of MS closely related to inflammation and changes in cytokines (Montoya et al., 2017; Ormstad et al., 2020; Patejdl, Penner, Noack, & Zettl, 2016), and it predominantly affects women (Faro et al., 2016). Excessive pro-inflammatory cytokine levels are causally associated with fever, fatigue, myalgias, sleep disturbances, and cognitive and mood disorders (Komaroff, 2017; Montoya et al., 2017) and they play a causal role in the onset of fatigue in MS (Ormstad et al., 2020). Despite the general acceptance of the cytokine hypothesis for MS-associated fatigue (Patejdl et al., 2016), the underlying processes of sex-specific fatigue-like symptoms remain poorly understood. MS symptoms such as fatigue can be triggered or exacerbated by elevated body temperature due to heat exposure, a phenomenon which has been coined Uhthoff’s phenomenon (Davis et al., 2008; Sumowski & Leavitt, 2014). Interestingly, even in the absence of changes in core temperature, heat intolerance or temperature sensitivity that only affects skin temperature may also contribute to this phenomenon (Christogianni et al., 2018). Accordingly, both fatigue and elevated temperature can be linked to aggravated systemic inflammation in MS (Heesen et al., 2006).

While thermal sensitivity represents a prominent symptom in MS, preclinical animal models commonly rely on clinical scoring of motor deficits with minimal attention to other types of biobehavioural endpoints. Alterations in body temperature and motor activity during EAE pathogenesis have been shown in male rats (Wrotek, Nowakowska, Caputa, & Kozak, 2020; Wrotek, Rosochowicz, Nowakowska, & Kozak, 2014). These findings indicate an association between altered thermoregulation (e.g., fever) and motor disturbances in EAE, however, these studies are limited to males. Although EAE disproportionally affects females, no EAE study has systematically evaluated sex differences in thermoregulatory responses and how these dynamics relate to EAE-induced symptoms.

The purpose of this study was to compare female and male C57BL/6 mice for susceptibility to EAE and their clinical progression using infrared (IR) thermography. IR thermography provides a non-invasive assessment of surface thermal changes in response to inflammation (Całkosiński et al., 2015) and stress (Faraji et al., 2021; Faraji & Metz, 2020; Tattersall, 2016) and accurately reflects variations in core temperature (Mei et al., 2018). Indeed, the peripheral autonomic nervous system that regulates perspiration and surface blood perfusion, determines heat patterns and gradients during aversive experiences and physiological challenges (Ioannou, Gallese, & Merla, 2014; Redaelli et al., 2019; Vinkers et al., 2013). Thus, cutaneous temperature variations may serve as a reliable physiological marker for psychobiological responses (Gjendal, Franco, Ottesen, Sørensen, & Olsson, 2018; Mei et al., 2018). Here, we report higher susceptibility among female mice to EAE based on greater clinical impairments, fatigue-like behaviours and pro- and anti-inflammatory cytokine levels. The neuroimmunological responses were accurately reflected by IR thermography of cutaneous temperature. These findings provide the first evidence of a sex-specific thermal response to immune-mediated demyelination, thus proposing a non-invasive assessment approach of the psychophysiological dynamics in EAE mice.

## Material and Methods

### Animals

Twenty-three adult male and female mice (C57BL/6 [B6]), 8–12 weeks old at the beginning of the study, were used. The animals were housed in pairs under a 12:12 h light/dark cycle with the light starting at 07:30 h. Animals were provided with water and food *ad libitum*. The room temperature was set at 22°C, and experimental procedures were conducted during the light phase of the cycle at the same time of day.

Animals were tested pre-induction and at three post-induction time points. The post-induction physiological and behavioural assessments began five days after day 0 of induction. The animals were also weighed daily and monitored carefully for any post-induction health impairments. Animals were euthanized once all IR imaging, behavioural testing and blood sampling were completed. The experimental design is illustrated in Figure 1**A**. All procedures were approved by the University of Lethbridge Animal Care Committee in compliance with the standards set out by the Canadian Council for Animal Care (CCAC).

**Figure 1:**
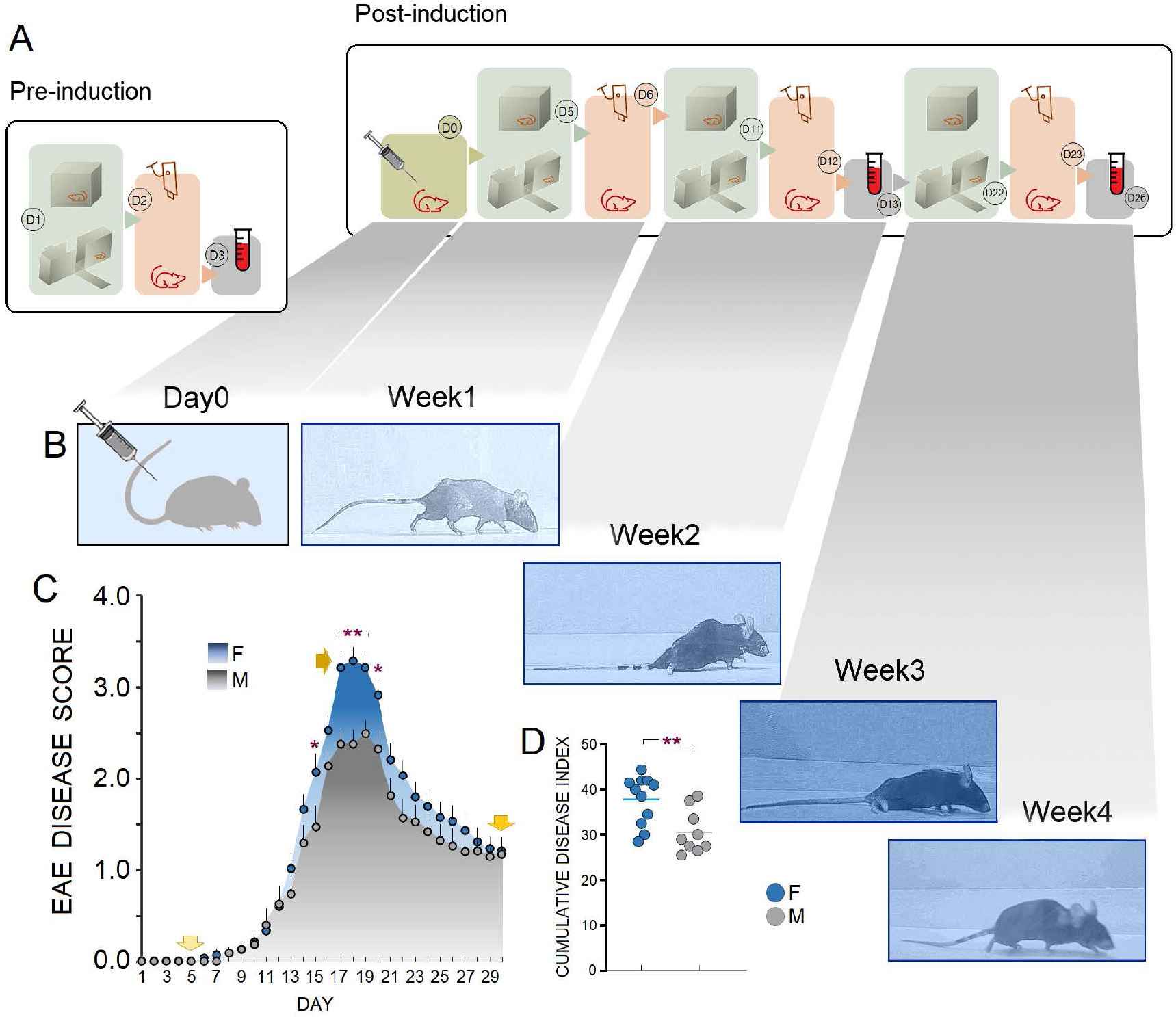
Experimental Design, and Clinical Severity and Progression of EAE. ***A***. *Schematic illustration of the time-course for the experiment*. ***B***. *Timeline illustrating clinical observations over a course of four weeks after induction of EAE (D0) in a representative female mouse. Week1*: although the tail was mobile, rigidity was felt in the tail while its tip was limp. *Week2*: limp tail and hind limb paralysis was detectable. In most animals, the hind limbs were not weight bearing and tail movement was absent. *Week3*: complete paralysis of hind limbs and tail. Forelimbs were used for ambulation. *Week4*: the tail was partially limp, and animals were able to regain weight support with hindlimbs. ***C***. *Disease progression after induction of EAE in female (n=11) and male mice (n=9)*. Yellow arrows indicate preclinical (D5), clinical or peak (D18), and chronic phases (D30) of EAE-induced symptoms in female mice. In both groups, the peak of symptoms occurred approximately between post-induction days 15-21. However, EAE in females revealed a significantly accelerated progression and aggravated symptom severity, particularly between days 17-19. ***D***. *Cumulative Disease Index (CDI) or total disease load generated through the summation of paralysis scores over time for each female and male animal*. Female EAE mice displayed a significantly higher CDI when compared with males. Horizontal bars indicate mean CDI in each group. Blue and grey circles represent individual animals (\**p* < 0.05; \*\**p* < 0.01, *O-W* ANOVA and independent *t*-test. Error bars show ± SEM).

### EAE Induction and Clinical Scoring

Prior to induction of EAE, baseline weights were recorded. Age-matched male and female mice were assigned to EAE (*n*=11-12/group) via right and left hind flank subcutaneous injection of 2 mg/ml of a mixture containing myelin oligodendrocyte glycoprotein MOG_35-55_ and *Mycobacterium tuberculosis* (strain *H37RA*) dissolved in phosphate-buffered saline (PBS) as described previously (Bettenson, Babatunde, Gustafson, Chan, & Metz, 2018). This mild form of EAE termed EAEnp did not involve the injection of pertussis toxin (PTx). Also, because EAE susceptibility is influenced by stress and handling procedures, the same experimenters (one male and one female) performed EAE induction and post-induction testing in all animals.

Following induction of EAE, animals were allowed to rest for 3 days. Post-induction weights were recorded daily to ensure weight loss did not exceed 25% of initial weight. Clinical scores were recorded in a blind manner using a 5-point clinical scoring scale (Hoghooghi et al., 2020). The clinical scoring scale followed predictable symptom progression from tail paralysis to quadriplegia and death (*0*, asymptomatic; *0*.*5-1*, partial to full tail paralysis; *2-2*.*5*, hind limb weakness to single hind limb paralysis; *3*, full hind limb paralysis; *4*, quadriplegia; *5*, moribund or death). Further, the summation of disease scores over time (cumulative disease index; CDI) was calculated from the daily clinical scores observed between day 6 and day 30 for mice in each group (Szalai, Hu, Adams, & Barnum, 2007).

### Infrared (IR) Imaging

The procedure for IR imaging was previously described by Faraji and Metz (2020). Briefly, animals (*n*=11-12/group) were placed individually in the center of a Plexiglas box (32 cm×32 cm) for 5 min per test session (pre-induction and post-induction days 6, 12, and 23). IR thermographic recording was performed with a FLIR IR thermographic camera (FLIR T450sc, Wilsonville, OR, USA; fixed emissivity=0.98 specified for skin in the manufacturer’s emissivity table) placed 80 cm above each animal to record dorsal temperature. The camera was able to follow changes in the animal’s surface temperature and its immediate surrounding with a thermal resolution of 320×240 pixels per image, thermal sensitivity of <30 mK at 30°C, and 60 Hz acquisition rate. IR imaging occurred in a windowless room with a steady temperature set at 22°C and relative humidity of ∼50%. Animals were protected from direct ventilation during IR imaging. Since body temperature fluctuates throughout the day, the time for IR thermal recording (8:00 am-10:30 am) was constant between animals and across time points. The post-induction IR thermal imaging was performed approximately one hour before behavioural assessments to reduce confounding effects of physical activity.

For the purpose of the thermal analysis, two oval-shaped principal regions of interest (ROI) were chosen. (1) *Head* (including eyes), covering a major portion of the frontal and parietal surfaces. An approximate measure of the sagittal suture allowed the elliptic ROI to split the top of the head into left and right sides. As well, a small segment of interparietal bone was included in either right or left ROIs. To control the effect of the position of each animal on the emitted thermal irradiations, the best postural condition for the head was chosen when rats were moving with their head oriented straight ahead without deviation to the side. (2) *Back*, the ROI included lower thoracic and upper and lower lumbar levels extended to the abdominal parts at equidistance from approximately 1.5 cm off the midline. In total, three ROIs (head [left and right] and back) were considered for analysis of changes in surface temperature.

IR thermal profiles were saved and analyzed using the FLIR image processing software (FLIR ResearchIR Max software 4.40.6.24). For sampling, 5 frames representing five time bins (one frame per each minute of the IR imaging) adjusted to the corresponding ROIs from the head and back were chosen for each animal. ROI sizes were identical for all frames and mice. The best-fit area to the ROIs in each frame/time bin was determined on the basis of the animal’s dorsal posture when approximately all relevant regions were bounded by the radius of the ellipses and/or when the animal was found in a prone position with all four limbs on the ground. The IR thermal analysis was performed by a male experimenter blind to the group identities.

### Behavioural Testing

#### Open Field

Open-field exploratory activity and emotional states were assessed using an AccuScan activity monitoring system (AccuScan Instruments Inc., OH, USA) with clear Plexiglas boxes [length 42 cm, width 42 cm, height 30 cm; (Ambeskovic et al., 2019)]. Animals (*n*=10-11/group) were placed individually into the open field and monitored for 10 min prior to induction of EAE (pre-induction) and at three time points after the induction (post-induction days 5, 11 and 22). The boxes attached to the computer recorded the activity based on sensor beam breaks. Total travel distance and margin travel distance (cm) were analyzed using VersaDatTM software (AccuScan Instruments Inc., OH, USA) to measure overall activity and emotional states, respectively. After each animal the apparatus was thoroughly cleaned with 10% Clinicide (Vetoquinol, Lavaltrie, QC, Canada) to eliminate any odor traces.

#### Elevated Plus Maze (EPM)

Anxiety-like behaviour was assessed using an EPM made of black Plexiglas. The apparatus consisted of two open and two closed arms (all 40 long × 10 cm wide) and was elevated 90 cm above the floor. The open arms had no side or end walls, whereas the closed arms featured side and end walls (40 cm high). Mice (*n*=7-8/group) were placed individually in the central square (10 cm × 10 cm) facing either the left or right open arm and were allowed to explore the maze for 5 min while being video-recorded. The standard measures (total time spent in open and closed arms) of the EPM were evaluated by a female experimenter blind to the experimental groups. In order to minimize olfactory cues, after each animal the apparatus was cleaned with 10% Clinicide [Vetoquinol, Lavaltrie, QC, Canada; (Faraji et al., 2018)] with modification).

#### Blood Sample Collection and Cytokine Assay

Blood samples for cytokine analysis were obtained on days without behavioural testing. According to previous procedures (Faraji & Metz, 2020), blood samples were taken a week before EAE induction (pre-induction) and at two post-induction time points (day 13 and immediately prior to euthanization on day 26). Mice were restrained by grasping the loose skin over the shoulder and behind the ears. A small puncture was made in the sub-mandibular vein using a lancet. Approximately 0.1 ml of blood was collected with a micro-tube. All samples were collected in the morning hours between 9:00 and 11:00 AM. Animals were returned to their home cage and allowed for recovery for 4 days. Plasma was obtained by centrifugation at 5,000 rpm for 10 min and stored at −80°C. A mouse Cytokine 23-Plex assay (Bio-Rad Laboratories, Montreal, QC, Canada) panel assessing pro- and anti-inflammatory cytokines (IL-2, IL-4, IL-5, IL-6, IL-7, IL-10, IL-12p70, IFN-γ, and TNF was conducted on a Luminex Bio-Plex Platform (Bio-Rad Laboratories, Montreal, QC, Canada). Peripheral plasma cytokine levels were determined using Bio-Rad Pro mouse cytokine multiplex commercial kits (Bio-Rad, Montreal, QC, Canada). Also, spleen and adrenal glands were weighed after euthanization to determine the impact of EAE on organ sizes. Since there were no differences between the right and left adrenal glands, an average of weights for both sides in each animal was considered for analysis.

#### Statistical Analysis

All data were analyzed with an SPSS 21.0 package (IBM SPSS Statistics, SPSS Inc., USA) and results are presented as mean ± SEM. Effects of main factors [Group—two levels; Time point—four levels; Time Bin—five levels) were analyzed as independent variables for the surface temperature in different ROIs, total and margin distance, total time in closed and open arms, and spleen and adrenal gland weights as dependent variables by *repeated measure (R-M) and one-way (O-W)* ANOVA. We also used a Linear Mixed-Model (LMM) approach to analyze the main and interaction effects of group (females and males) and sampling time points (pre-induction and post-induction days 13 and 26) for the changes in circulating cytokine levels. The LMM, contrary to the simple R-M ANOVA, integrates the data available for all subjects while adjusting estimates for any missing data within groups. The LMM was also followed by separate O-W ANOVA to test for the effect of Group on every single time point. The experimental design allowed us to also pursue separate means comparisons between groups using independent sample *t*-tests for within-subject comparisons. To evaluate the magnitudes of the effects, effect sizes (*η*^2^ for ANOVA and Hedges’ *g* for *t*-tests) were calculated. Values of *η*^2^ = 0.14, 0.06, and 0.01 were considered for large, medium, and small effects, respectively. Also values of *g* = 0.8, 0.5, and 0.2 were considered for large, medium, and small effects in Hedges’ *g*, respectively. A *p*-value of less than 0.05 was considered statistically significant.

## Results

### EAE Clinical Score: *Disease Severity in EAE is Sex-Dependent*

EAE disease progression is shown in Figure1**B**. Clinical signs in males and females ranged from tail weakness in the mild cases to full hindlimb paralysis. The peak of clinical disabilities in both groups occurred between days 15-21 post-induction. At the peak of disability, animals in each group exhibited hindlimb impairment or paralysis. However, female EAE mice (*n*=11) displayed a tendency for higher clinical scores than males (*n*=9; 1.25±0.05 vs. 1.02±0.05). A main effect of Group (*F*_1,18_=9.41, *p*<0.007, η^2^=0.34) with no interaction between Group and Time (*p*>0.05) was approved by *R-M* ANOVA (Figure1**C**). Also, a separate between-group comparison indicated a significantly accelerated disease progression in females based on days 15 (*p*<0.04), 17 (*p*<0.01), 18 (*p*<0.01), 19 (*p*<0.01), and 20 (*p*<0.05) post-induction (all *O-W* ANOVA). The CDI or the measure of disease severity calculated for days 6-30 post-induction also indicated more severe disease in females than males (CDI: 37.72±1.63 vs. 30.61±1.59; *t*_18_= 3.06, p<0.01, *g*= 1.38, independent samples *t*-test; Figure1**D**). Thus, both clinical scores at peak time points and the overall course of EAE were significantly exacerbated in females compared to males.

### IR Cutaneous Temperature: *Sex Differences Regulate Thermal Responses in EAE*

Figure 2**A** shows the procedure of IR imaging accompanied by a representative thermal image of surface temperature (*right panel*) taken from two regions of interest in head (left and right) and back. Because there were no differences in the thermal response between the left and right sides of the head, and no effect of Time Bin, the averages of left and right head cutaneous temperatures were used for analysis. Females (*n*=11) and males (*n*=12) displayed a similar pattern in cutaneous temperatures across all five time bins in head and back. Analysis of the mean surface temperature of the head and back for each group in the pre-induction session and day 6 post-induction by *O-W* ANOVA did not show significant effects of Group [(female vs. male, *Pre-induction (head)*: 36.6±0.1°C vs. 36.3±0.1°C, *(back)*: 35.6±0.1°C vs. 35.7±0.1°C; *Post-induction day6 (head)*: 37.0±0.1°C vs. 36.8±0.1°C, *(back)*: 36.3±0.2°C vs. 36.0±0.1°C, all *p*>0.05; Figure 2**B**&**C**)]. However, a significant effect of Group (*F*_1,21_=25.98, *p*<0.001, *O-W* ANOVA) on post-induction day 12 indicated that the heat emitted from head in females in response to EAE induction was greater than males (37.2±0.1°C vs. 36.4±0.1°C; Figure 2**D**). Females and males were not different in the surface temperature at the back on day 12 post-induction (*p*>0.93). Moreover, the average temperature for both head and back in females was significantly higher than males on day 23 post-induction (*head*: 37.8±0.1°C vs. 36.4±0.1°C; *F*_1,21_=50.64, *p*<0.001, *back*: 37.4±0.1°C vs. 36.6±0.1°C; *F*_1,21_=8.92, *p*<0.01, *O-W* ANOVA; Figure 2**E**). Also, when thermal responses to EAE at the head and back were analyzed, females displayed a completely different profile of thermal changes than males across all four time points (Figure 2**F**-**K**). *R-M* ANOVA indicated females emitted significantly more heat from the head than the back (effect of Region: *F*_1,20_=25.4, *p*<0.001, η^2^=0.56). No interaction, however, was found between Region and Time point (*p*>0.25). The significant effect of Region in females disappeared on day 23 post-induction where both regions responded to EAE similarly (Figure 2**G**). Also, rate of change (ROC) during all four time points showed that females experienced greater thermal changes at the back than the head from pre-induction to post-induction day 23, although in an increasing pattern (Figure 2**H**). In contrast, changes of thermal profiles in males followed a different pattern of surface temperature in head and back. No significant effect of Region (*p*>0.05) or interaction between Region and Time point (*p*<0.001) were found in males. However, a further *O-W* ANOVA indicated that heat emitted from head pre-induction (36.3±0.1°C vs. 35.7±0.1°C; *F*_1,22_=9.81, *p*<0.01, *O-W* ANOVA) and on post-induction day 6 (36.8±0.1°C vs. 36.0±0.1°C; *F*_1,22_=14.34, *p*<0.01, *O-W* ANOVA; Figure 2**J**) was significantly higher than back, whereas the between-region differences disappeared on days 12 and 23 (All *p*>0.05) with a slightly enhanced thermal response at the back at both time points. The ROC analysis also indicated that, unlike females, the impact of EAE on surface temperature on head and back was divergent in males; the back responded to EAE with elevated temperature whereas head emitted lower heat, particularly at chronic time points (Figure 2**K**).

**Figure 2:**
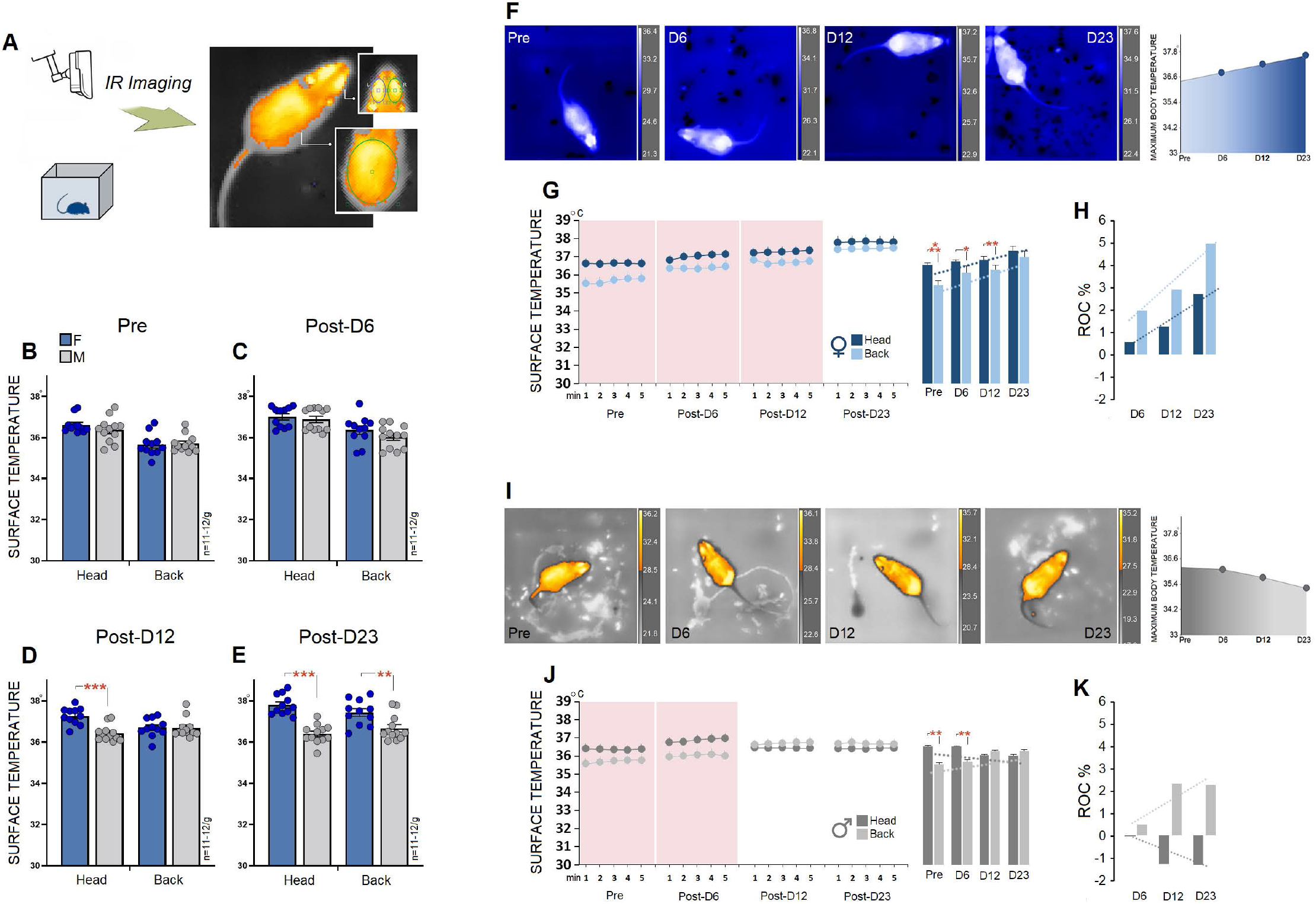
Infrared Thermography of Surface Temperature. ***A***. *Setup*. An infrared (IR) imaging camera was placed 80 cm above the animal and followed changes in the animal’s surface temperature in real time. Also, thermal images (*right panel*) show two regions of interest [head (left and right) and back in assessments of surface temperature]. ***B-E***. *Sex-specific thermal responses to EAE in different regions of interest and time points*. Females and males showed a differential temporal profile of thermal response to EAE in the head and back. Blue and grey circles in panels represent individual animals. ***F***. *Representative dorsal view plots of the maximum surface temperature of a freely moving female mouse at different time points*. Temperatures (°C) are indicated by the color code in the right panels. The animals displayed an increasing thermal changes in the body from pre-induction (36.4°C) to post-induction day 23 (37.6°C). ***G***. *Comparison of cutaneous thermal changes in head and back of females*. Females responded to EAE with enhanced surface temperatures in head and back on all post-induction days, with the largest change on day 23. ***H***. *The rate of changes (ROC)*. Females experienced greater post-induction cutaneous thermal changes in the back than head. However, the regional thermal responses to EAE in females in both ROIs follow a similar temporal profile. ***I***. *Four representative dorsal view plots of the average body surface temperature of a freely moving male mouse at different time points*. As shown by variations during four experimental conditions that are depicted by the color code plots in the right panels, the mouse displayed a reduced thermal response from 36.2°C in the pre-induction phase to 35.2°C on day 23 post-induction. ***J***. *Changes in cutaneous thermal responses in males across four time points of IR imaging*. Unlike females, males responded to EAE induction with reduced thermal changes in the head and an increased thermal response in the back. ***K***. *The rate of changes (ROC) in thermal responses in males*. Note, the trendlines of thermal changes in the head and back in males illustrate sex-specific differences. Surface temperature in the head and back of males diverged in the back and head during post-induction days. (*n*=11-12/g, \*\**p* < 0.01, \*\*\**p* < 0.001, *O-W* ANOVA). Post-D6, Post-D13 and Post-D23 indicate post-induction days.

In summary, both sexes experienced remarkable degrees of cutaneous thermal alterations in both ROIs in response to induction of EAE. Also, females were more vulnerable to EAE than males as shown by increased surface temperature in the head and back. Even though females and males responded similarly in head and back temperatures on post-induction day 6, the ROC across experimental sessions indicated that patterns in females appeared to follow an order of (head>back*) < (head>back*) < (head>back*) < (head>back) which was different from the order of (head>back*) < (head>back*) > (head<back) > (head<back) observed in males (* denotes significant difference).

### Locomotion and Emotionality: *Sex-Dependent Psychomotor Inhibition and Fatigue in EAE*

An open field activity monitoring system (Figure3***a***-***c***-top) was used to assess exploratory and emotional behaviours based on overall activity (total distance) and path taken by animals in the margin area at four time points (pre-induction, post-induction days 5, 11, and 22). No significant main effect of Group was found for total distance (*p*>0.12, *R-M* ANOVA). However, females (*n*=9) traveled significantly less than males (*n*=11) on the chronic time point (post-D22, 140.77±13.47 cm vs. 203.09 ±13.91 cm, *F*_1,19_=10.06, *p*<0.01, *O-W* ANOVA; Figure3**D**-top). No significant group difference in total distance on post-induction day 11 was found despite slightly reduced distance in females (154.33±15.71 cm vs. 192.81±11.36 cm, *p*>0.057; *O-W* ANOVA). There was no main effect of Group (*p*>0.40, *R-M* ANOVA), however, Time point (*F*_3,54_=3.03, *p*<0.03, η^2^=0.14, *R-M* ANOVA) in terms of margin distance or path taken near walls indicate that margin distance only on post-induction days 11 and 22 was impacted by EAE. An *O-W* ANOVA revealed that females traveled significantly less in the margin area than males on post-induction day 22 (171.63±12.19 cm vs. 132.55±12.99 cm, *F*_1,18_=4.77, *p*<0.04, *O-W* ANOVA; Figure3**E**-top). Thus, EAE was associated with impaired locomotion in females particularly at chronic time points.

**Figure 3.**
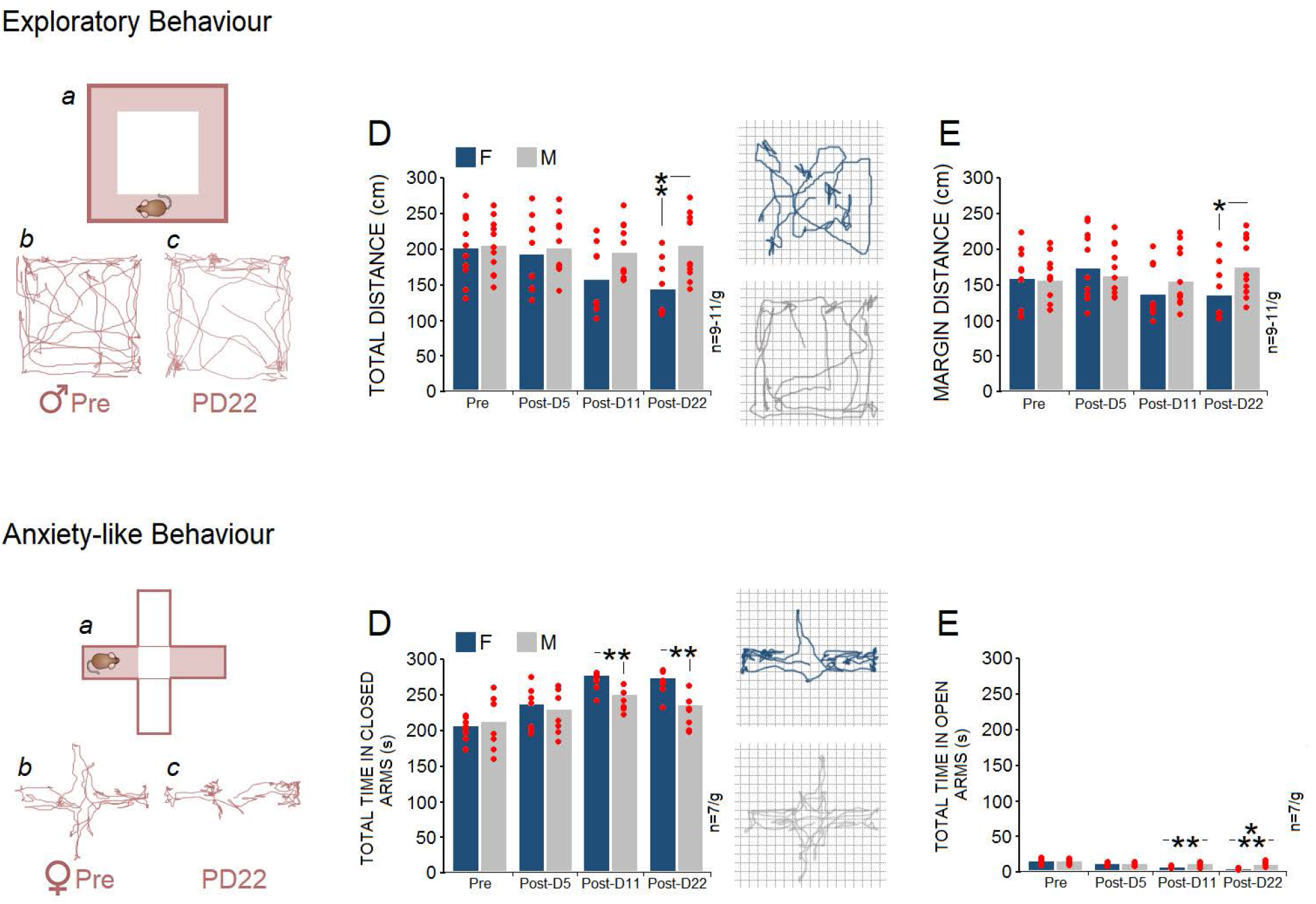
Behavioural Profiles of Fatigue. ***a-c***-top. *Representative exploratory locomotor trajectories of a male mouse in the open field pre- and post-induction*. Darker spots display stops and/or time spent in corners near the walls. ***D*** & ***E***. *Total and margin distance during open field exploration*. Path length or distance traveled by females (*n*=9) and males (*n*=11) indicated that females traveled significantly less distance mainly in the central area than males on post-induction day 22. ***a-c***-bottom. *Representative paths traveled by a female mouse in the elevated plus maze pre- and post-induction*. ***D*** & ***E***-bottom. *Total time spent in closed and open arms by females and males (n=7/g)*. Although EAE increased the time spent in the closed arms in both sexes, females spent more time in the closed arms and less in the open arms compared to males on post-induction days 11 and 22. Red circles represent individual animals (\**p* < 0.05; \*\**p* < 0.01, \*\*\**p* < 0.001, *O-W* ANOVA). Post-D5, Post-D11 and Post-D22 indicate post-induction days.

Anxiety-related behaviour was also examined using the EPM (Figure3***a-c***-bottom). A significant main effect of Group (*F*_1,12_=6.84, *p*<0.02, η^2^=0.36, *R-M* ANOVA) and Time point (*n*=7/g, *F*_3,36_=14.19, *p*<0.01, η^2^=0.81, *R-M* ANOVA) with no interaction (*p*>0.43) was found in total time spent in the closed arms. A separate *O-W* ANOVA conducted to compare groups at each time point showed that females spent more time than males in the closed arms on post-induction day 11 (274.14±5.22 s vs. 247.14±5.92 s, *F*_1,12_=11.68, *p*<0.01) and day 22 (269.42±6.74 s vs. 232±9.10 s, *F*_1,12_=10.91, *p*<0.01; Figure3**D**-bottom), suggesting that females displayed increased anxiety-related behaviours. Total time spent in open arms also provided further evidence of anxiety in females. Again, both main effects of Group (*n*=7/g, *F*_1,12_=14.52, *p*<0.01, η^2^=0.54, *R-M* ANOVA) and Time point (*F*_3,36_=25.35, *p*<0.001, η^2^=0.67, *R-M* ANOVA) as well as the interaction between Group and Time point (*F*_3,36_=5.38, *p*<0.01, η^2^=0.31, *R-M* ANOVA) were significant. A separate *O-W* ANOVA conducted for total time in open arms indicated that females spent less time in open arms than males on post-induction day 11 (4.14±0.45 s vs. 9.57±1.37 s, *F*_1,12_=13.97, *p*<0.01) and day 22 (1.28±0.52 s vs. 7.42±1.25 s, *F*_1,12_=20.54, *p*<0.001; Figure3**E**-bottom). These observations indicate a much greater anxiety-like response to EAE in females than males, particularly at chronic time points when there was no sex difference in terms of clinical disease progression.

### EAE-Related Immunological Markers: *Spleen and Adrenal Gland Size, and Total Cytokine Profiles Reveal Sex Differences*

Spleen (*t*_16_=3.38, p<0.01, *g*=0.53, independent samples *t*-test) and adrenal glands (*t*_17_=2.16, p<0.04, *g*=0.37, independent samples *t*-test) weights (Figure4**A**) revealed a sex-dependent effect where both organs in females weighed more than in males (*spleen*: .103±.008g vs. .069±.004g; *adrenal gland*: .005±.0002g vs. .004±.0002g). The enlarged spleen and adrenal glands may indicate inflammatory deficiency and chronic stress in females, respectively.

**Figure 4.**
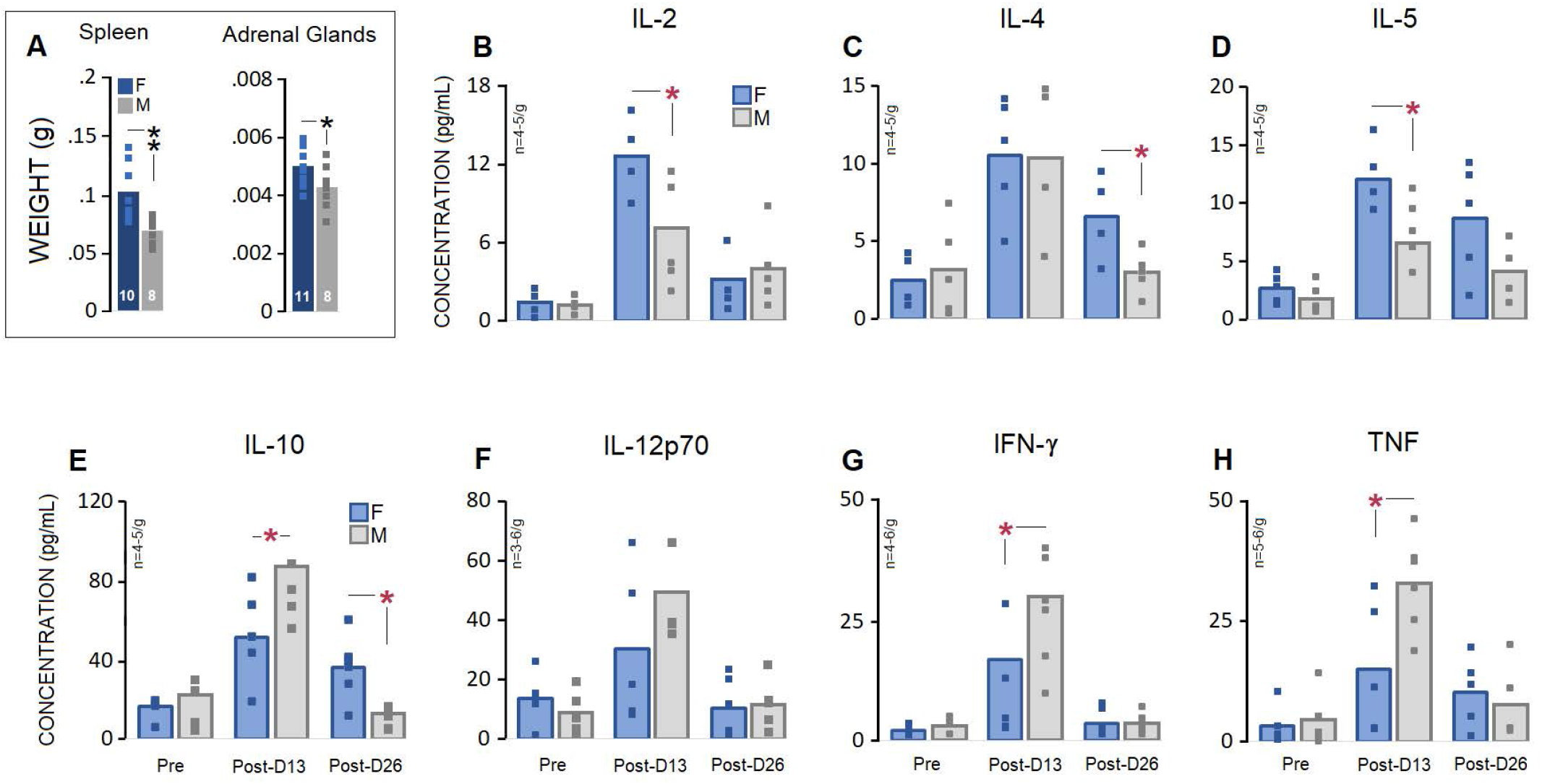
Immunological Markers Induced by EAE. ***A***. *Sex-specific changes in spleen and adrenal gland sizes in response to EAE*. Both spleen and adrenal gland weights significantly increased in females (*n*=10-11) compared with males (*n*=8). ***B****-****H***. *Sex-specific profiles of pro- and anti-inflammatory cytokine responses in EAE*. Plasma cytokine levels across three time points (pre-induction, and post-induction days 13 and 26) showed upregulated IL-2 and IL-5 on PD13, and IL-4 and IL-10 on post-induction day 26 in females. Also, females exhibited downregulation of IL-10, IFN-γ, and TNF on PD13. Blue and grey squares in the graphs represent individual animals in each group (\**p* < 0.05, *O-W* ANOVA). Post-D13 and Post-D26 indicate post-induction days.

Pro- and anti-inflammatory responses to EAE were measured via changes in circulating plasma cytokine concentrations across three time points (pre-induction, post-induction days 12 and 26; Figure4**B**-**H**).

*Interleukin-2 (IL-2)*. The linear mixed-model (LMM) showed no main effect of Group (*p*>0.08) but Time point (*F*_2,22_=22.98, *p*<0.001, *type III tests of effects*). Follow-up *O-W* ANOVA indicated that females displayed higher IL-2 concentration than males at post-induction day 13 (12.63±1.54pg/mL *vs*. 6.51±1.86 pg/mL, *F*_1,7_=5.94, *p*<0.04, *n*=4-5/g; Figure4**B**).

*Interleukin-4 (IL-4)*. No main effect of Group (*p*>0.41) but Time point (*F*_2,21_=13.70, *p*<0.001, *LMM*-*type III tests of fixed effects*) was observed in IL-4 concentration. A separate analysis conducted by *O-W* ANOVA confirmed the results of *LMM* analysis where females displayed higher IL-4 levels than males on post-induction day 26 (6.53±1.37pg/mL *vs*. 3.01±.58 pg/mL, *F*_1,7_=6.52, *p*<0.03, *n*=4-5/g; Figure4**C**).

*Interleukin-5 (IL-5)*. Both groups showed similar responses to EAE with higher levels of IL-5 on post-induction days 13 and 26. LMM analysis indicated a main effect of Group (*F*_1,22_=9.49, *p*<0.01, *Type III tests of fixed effects*) and Time point (*F*_2,22_=17.50, *p*<0.001, *Type III tests of fixed effects*). Follow-up analysis by *O-W* ANOVA, however, indicated that only females displayed greater concentration of IL-5 than males on post-induction day 13 (12.53±1.49 pg/mL *vs*. 7.76±1.25 pg/mL, *F*_1,7_=6.04, *p*<0.04, *n*=4-5/g; Figure4**D**).

*Interleukin-10 (IL-10)*. Induction of EAE elicited a IL-10 response in both females and males. However, females showed lower concentration of IL-10 on post-induction day 13 and higher IL-10 levels on day 26 compared to males. *LMM* revealed no main effect of Group (*p*>0.33) but Time point (*F*_2,22_=27.44, *p*<0.001, *Type III tests of effects*). Moreover, *O-W* ANOVA confirmed that the IL-10 level in females was significantly different from males at both post-induction time points (*PD13*: 52.43±10.86 pg/mL *vs*. 88.64±8.55 pg/mL, *F*_1,7_=6.29, *p*<0.04; *PD26*: 36.47±8.10 pg/mL *vs*. 13.04±2.82 pg/mL, *F*_1,8_=7.63, *p*<0.02, *n*=4-5/g; Figure4**E**).

*Interleukin-12p70 (IL-12)*. Females and males responded to EAE with similar raises in IL-12 levels. The *LMM* showed no effect of Group (*p*>0.36) but Time point (*F*_2,22_=10.73, *p*<0.001, *Type III tests of effects*). Nevertheless, there were no significant differences between groups on post-induction days 13 and 26 (all *p*>0.05, *O-W* ANOVA, *n*=3-6/g; Figure4**F**).

*Interferon gamma (IFN-γ)*. There was a main effect of Group (*F*_2,25_=5.28, *p*<0.03, *LMM-type III tests of effects*) and Time point (*F*_2,25_=16.97, *p*<0.001, *LMM-type III tests of effects*) where females displayed a reduced level of IFN-γ on post-induction day 13 compared to males (10.70±5.12 pg/mL *vs*. 27.94±4.93 pg/mL, *F*_1,9_=5.80, *p*<0.03; *O-W* ANOVA, *n*=4-6/g; Figure4**G**).

*Tumor Necrosis Factor (TNF)*. Females responded to EAE differently than males, particularly on post-induction day 13. Again, the *LMM* analysis indicated no main effect of Group (*p*>0.09) but Time point (*F*_2,25_=15.14, *p*<0.001, *Type III tests of effects*). When compared with males, however, females displayed lower concentration of TNF on post-induction day 13 (15.55±6.35 pg/mL *vs*. 33.79±4.12 pg/mL, *F*_1,9_=6.19, *p*<0.03; *O-W* ANOVA, *n*=5-6/g; Figure4**H**).

## Discussion

The contribution of thermoregulatory activity to EAE-associated symptoms, which are potentially relevant to human MS, is unknown, but may in part explain the sexual dimorphisms found in both conditions. Based on our previous observation of sex-specific thermoregulatory activity in response to stress (Faraji & Metz, 2020), the goal of this study was to determine if sexual dimorphisms in thermoregulatory dynamics are associated with the progression of EAE. Here, using IR imaging, we show that: (1) female C57BL/6 mice display higher susceptibility than males to the clinical course and severity of a mild form of EAE that may lead to (2) their greater thermal responses to EAE during acute and chronic phases, (3) disruption of locomotor activity and emotionality, and (4) distinct inflammatory responses. These findings are the first to identify a sexually dimorphic profile of thermoregulation linked to EAE pathogenesis.

The C57BL/6 model of EAE generates a characteristic acute monophasic profile and stable chronic disease course (Berard et al., 2010). However, despite the clear female-biased differences in MS incidence and outcome, the C57BL/6 females’ preponderance in EAE susceptibility is still controversial (Hoghooghi et al., 2020; Ondek, Nasirishargh, Dayton, Nuño, & Cruz-Orengo, 2021; Papenfuss et al., 2004). In the present study, we observed a sexual dimorphism in C57BL/6 mice with respect to disease outcomes. Similarly to the SJL/J strain (Voskuhl, Pitchekian-Halabi, MacKenzie-Graham, McFarland, & Raine, 1996), C57BL/6 female mice also showed a more severe disease phenotype than males, reflected by greater neurological deficits, in a model of mild EAE. Both clinical scores at peak time points and the overall course of EAE were significantly exacerbated in females compared to males, a finding in accordance with the previous literature (Hoghooghi et al., 2020). Any potential discrepancy observed between the present results and earlier investigations that have found no sex differences in the C57BL/6 mouse model of EAE might derive from age, the EAE induction regimen and experimental demands, the severity of the disease course, as well as procedural stress induced by the impact of animal handling, testing or the experimenter’s sex. Confounding effects of the experimenter’s sex may play a critical role in causing sex-biased differences in rodent models. Indeed, the opposite-gender dynamics that have been recently reviewed in humans (Chapman, Benedict, & Schiöth, 2018) may be reproduced in preclinical studies (Faraji et al., 2021; Sorge et al., 2014) and are typically linked to psychophysiological distress. Accordingly, stress-induced hyperthermia and infection-induced fever are two distinct physiological processes that need to be differentiated by comprehensive biobehavioural testing (Harden, Kent, Pittman, & Roth, 2015; Spencer, Heida, & Pittman, 2005; Vinkers et al., 2009).

Stressed animals may experience high emotionality mediated by the activity of the hypothalamic-pituitary-adrenal (HPA) axis that can modulate physiological and behavioural responses to disease pathogenesis. Notably, a central sex-specific mechanism involved in stress regulation highlights the key role of the HPA axis in diverging stress responses in females and males (McCarthy, Arnold, Ball, Blaustein, & De Vries, 2012; Oyola & Handa, 2017), thus generating distinct neurohormonal stress responses in females compared to males. Moreover, the profile of HPA axis activity is characterized by prominent sex differences. In fact, females initiate the HPA-axis activity more rapidly in response to emotional stimuli and produce a greater output of stress hormones (Heck & Handa, 2019; Kokras, Hodes, Bangasser, & Dalla, 2019). This may also explain, at least partially, the greater adrenal weight in females seen in the present study.

According to the present findings, the sex-dependent clinical symptoms and cytokine profiles of EAE were reflected by altered thermoregulatory activity. While thermal responses to EAE in rodents have not been extensively studied, it is widely accepted that hormonal influences, such as estrogens, may modulate central subsystems that determine cutaneous and subcutaneous thermal responses (Charkoudian, Hart, Barnes, & Joyner, 2017; Charkoudian & Stachenfeld, 2014). The thermal maps of both elevated head and back temperature in EAE post-induction phases among females may portray sex-specific cutaneous vascular changes, which is contrasted by a marked drop in head temperature versus an increase in back temperature among males. In accord with our previous report (Faraji & Metz, 2020), this finding shows that the surface thermal maps in females are rather generalized and not regionally restricted to a particular area of the body. Whether the enhanced temperature represents a low-magnitude sympathetic vasoconstrictor signal to the skin arteries that facilitates skin blood flow (Blessing, 2003) or more general central processes that induce a passive thermal change through blood circulation (Lim, Byrne, & Lee, 2008), is beyond the scope of the present study. Nevertheless, these changes reveal the sex-specific nature of thermoregulatory activity in mice in response to homeostatic challenges.

Despite the lack of a definitive febrile response state, the present findings cannot rule out that the cutaneous temperature in EAE females reflects a transient fever following induced inflammation. However, because thermal responses in the present study were measured at room temperature (22°C) below the thermoneutral zone for rats, any statement about infection-induced fever should be made with caution. It was also suggested that heat-related fatigue in human MS arguably represents a form of central fatigue that reflects lack of energy that, in contrast to peripheral factors, fails to sustain drive to spinal motoneurons (Davis, Wilson, White, & Frohman, 2010; Gandevia, Allen, Butler, & Taylor, 1996). Therefore, the thermoregulatory dysfunction in MS and increased symptomatology related to fatigue are likely due to impairments in central conduction, specifically in the hypothalamus (Davis et al., 2010). Moreover, MS is accompanied by leukocyte infiltration to the central nervous system (CNS), which coincides with production of cytokines and vasoactive substances that in turn affect demyelination and neuronal degeneration (Göbel, Ruck, & Meuth, 2018). A thermoregulatory response to cytokine activation appears to be a disease tolerance defense strategy (Schieber & Ayres, 2016), while both pro- and anti-inflammatory cytokine release as correlates of thermoregulation act in a sex-dependent manner (Menees et al., 2021; Russi, Ebel, Yang, & Brown, 2018; van den Broek, Damoiseaux, De Baets, & Hupperts, 2005; Zhang et al., 2012) arguably via the influence of sex hormones (Klein & Flanagan, 2016). The significant splenomegaly in females may be indicative of modifications in the number and distribution of immune cells within the spleen (Menees et al., 2021) and/or more myeloid-derived suppressor cells (MDSCs) in the spleen that lead to a greater infiltration of the CNS (Melero-Jerez et al., 2020) in females.

Consistent with the behavioural and physical findings, the plasma values for immune markers indicated a sex-specific pattern of activity in 7 cytokines (IL-2, IL-4, IL-5, IL-10, IL-12p70, IFN-γ, and TNF) following EAE induction. Here, both sexes displayed an augmented production of cytokines at different time points, especially in the acute phase of EAE (post-induction day 13). We show, however, that the nature of cytokine production and signaling in EAE differs between females and males, a finding that may explain symptomatic aggravation in females with EAE or MS. For example, increased interleukin-2 (IL-2) levels in females may explain their greater disease severity because the IL-2-induced cytokine cascade can result in early blood brain barrier impairment and focal vascular edema, the hallmarks of MS lesions (Gallo et al., 1992). Furthermore, IL-2 levels in MS patients are positively correlated with clinical scores (Sharief & Thompson, 1993). Because IL-2 levels are increased in MS patients and because IL-2 mediates T cell proliferation and differentiation into effector cells, it can be valuable to assess how sex influences early inflammatory responses in T cell-mediated immune disorders such as MS. It appears that females have normally a higher IL-2 release compared to males (Gilli, DiSano, & Pachner, 2020). However, because our measure of IL-2 in pre-EAE showed no sex differences, the higher level of IL-2 in females post-induction may also be explained as a direct sex-specific early response to the EAE pathogenesis.

The present data suggest a female-biased deficiency to produce IL-10. IL-10 production by innate immune cells, which is also regulated by androgens, is believed to be a protective anti-inflammatory cytokine in EAE (Dai, Ciric, Zhang, & Rostami, 2012; Fletcher, Lalor, Sweeney, Tubridy, & Mills, 2010). Further, it is known that IL-10 secretion is decreased in MS patients (Göbel et al., 2018) and low levels of IL-10 are a predictor of MS-related symptoms including demyelinating events (Wei et al., 2019). The significant up- and down-regulation of IL-10 at acute and chronic time points (post-induction days 13 and 26, respectively) appears contradictory to the normal profile of IL-10 in males (Gilli et al., 2020). Thus, IL-10 production and sex differences may vary across disease stages of EAE. The impaired IL-10 signaling in females may enhance inflammatory response that potentially results in exacerbated immunopathology and/or tissue damage.

Another finding involved lower levels of interferon gamma (IFN-γ) and tumor necrosis factor alpha (TNF) in females compared with males on post-induction day 12. Because myelin-specific Th1 cells, which play a causal role in EAE induction (Voskuhl & Palaszynski, 2001), produce IFN-γ and TNF (Göbel et al., 2018), this finding provides further evidence that deficiency of these two multifunctional cytokines can lead to an early sex-based impaired response to EAE and a more severe clinical EAE phenotype in females. Nevertheless, one should be cautious to link such alterations in cytokines to EAE pathogenesis because changes in circulating cytokine profiles may reflect either cause or effect, or a combination of the two, in the female preponderance for EAE susceptibility.

Importantly, the sex bias in immune-inflammatory pathways may also lead to variations in psychological and behavioural responses to autoimmune diseases such as MS. Fatigue and depression that are clinically associated with anhedonia affect the majority of MS patients (Heitmann et al., 2020) and affect more women than men (Faro et al., 2016). In the present study, female mice responded to EAE induction with reduced locomotion and increased anxiety-like behaviour. Psychomotor inhibition displayed by females, mostly on chronic post-induction days, not only reflects greater vulnerability of females to sickness behaviours, but also indicates a sex-biased profile of motivational and emotional defects in EAE that develops independently from early impairments in motor function (Acharjee et al., 2013; Gold & Irwin, 2009). Furthermore, because fatigue-like behaviours in the present study occurred in association with cytokines such as IL-4 and IL-10, fatigue-like behaviours may be causally linked to cytokine-mediated pathogenesis and brain inflammatory cell infiltration. Although biologically heterogenous, the impact of cytokines on behaviour (Pedraz-Petrozzi, Neumann, & Sammer, 2020) appears to follow a female-biased trajectory in EAE.

Interestingly, endogenously elevated body temperature without heat exposure is linked to fatigue in MS patients (Sumowski & Leavitt, 2014), a phenomenon which has not been described yet in an animal model. Here, we showed enhanced thermoregulatory activity in response to EAE on post-induction day 23 in association with impaired locomotion in females. In contrast to males, females also exhibited a generalized heat emission that progressively extended to both head and back in response to EAE. In clinical populations including MS patients, functional decrements typically shown by fatigue may begin to appear when temperature exceeds normal levels, even in a range of 0.5 to 1.0°C (Guthrie & Nelson, 1995; White, Vanhaitsma, Vener, & Davis, 2013). Even though MS-related symptoms cannot be directly reproduced in an induced animal model, reduced exploratory activity and impaired response to novelty in open field and elevated plus mazes offer a translational concept of fatigue in MS (Grace et al., 2017). As fatigue predominantly affects female patients with MS, the present findings confirm a sex-dependent thermoregulatory mechanism linked to neurological impairments in EAE. Thus, the present findings provide further support for the validity of EAE as a model of human MS and offers new insights into the link of motor and non-motor symptomatology in this condition.

## Acknowledgements

The authors are very grateful to Dr. Isabelle Gauthier for support in EAE model development, and to Dr. Quentin Pittman for thoughtful comments on the manuscript. The authors acknowledge support by the Alberta Innovates-Health Solution Collaborative Research and Innovation Opportunities (CRIO) program (VY, GM), Natural Sciences and Engineering Research Council (NSERC) of Canada Discovery Grants #5628 and #31 (GM), and Canadian Institutes of Health Research (CIHR) Project Scheme #363195 (GM). The funding agencies had no role in the design of the study, collection and analysis of data, the decision to publish or in writing of the manuscript.

